# Feammox *Acidimicrobiaceae* bacterium A6, a lithoautotrophic electrode-colonizing bacterium

**DOI:** 10.1101/300731

**Authors:** Melany Ruiz-Urigüen, Weitao Shuai, Peter R. Jaffé

## Abstract

An *Acidimicrobiaceae* bacterium A6 (A6), from the Acitnobacteria phylum was recently identified as a microorganism that can carry out anaerobic ammonium oxidation coupled to iron reduction, a process also known as Feammox. Being an iron-reducing bacterium, A6 was studied as a potential electrode-reducing bacterium that may transfer electrons extracellularly onto electrodes while gaining energy from ammonium oxidation. Actinobacteria species have been overlooked as electrogenic bacteria, and the importance of lithoautotrophic iron-reducers as electrode-reducing bacteria at anodes has not been addressed. By installing electrodes in soil of a forested riparian wetland where A6 thrives, as well as in A6 bioaugmented constructed wetland (CW) mesocosms, characteristics and performances of this organism as an electrode-reducing bacterium candidate were investigated. In this study, we show that *Acidimicrobiaceae* bacterium A6 is a lithoautotrophic bacterium, capable of colonizing electrodes in the field as well as in CW mesocosoms, and that it appears to be an electrode-reducing bacterium since there was a boost in current production shortly after the CWs were seeded with *Acidimicrobiaceae* bacterium A6.

**IMPORTANCE:** Most studies on electrogenic microorganisms have focused on the most abundant heterotrophs, while other microorganisms also commonly present in electrode microbial communities such as Actinobacteria have been overlooked. The novel *Acidimicrobiaceae* bacterium A6 (Actinobacteria) is an iron-reducing bacterium that can colonize the surface of anodes and is linked to electrical current production, making it an electrode-reducing candidate. Furthermore, A6 can carry out anaerobic ammonium oxidation coupled to iron reduction, therefore, findings from this study open up the possibility of using electrodes instead of iron as electron acceptors as a mean to promote A6 to treat ammonium containing wastewater more efficiently. Altogether, this study expands our knowledge on electrogenic bacteria and opens up the possibility to develop Feammox based technologies coupled to bioelectric systems for the treatment NH_4_^+^ and other contaminants in anoxic systems.

## INTRODUCTION

Electrode-reducing bacteria (ERB) are part of a group of electrogenic microorganisms that have the ability to extract energy from different types of electron donors such as organic matter, and transfer those electrons to various terminal electron acceptors including electrodes operating as anodes, in which case a low-density electrical current is produced (1). Known electrogenic microorganisms include yeast and various bacteria (2). Studies on community composition analysis of ERB show ample taxonomic diversity mostly dominated by three phyla, Firmicutes, Acidobacteria, and Proteobacteria, the latter contains some of the most commonly present and extensively studied ERB: *Geobacter spp*. and *Shewanella spp*. (1–4). Most of these organisms are heterotrophs that thrive in anaerobic environments and obtain their energy by oxidizing organic matter (1). Commonly, ERB are iron-reducing bacteria (FeRB) (1), and many depend on or benefit from electron shuttles to facilitate the transfer of electrons from the microorganism to a solid electron acceptor such as the iron oxides [Fe(III)] (5).

*Acidimicrobiaceae* bacterium A6 (referred to as A6 from here on) is an autotrophic anaerobic microorganism that obtains its energy by oxidizing ammonium (NH_4_^+^) to nitrite (NO_2_^-^) and transferring the electrons to oxidized iron [Fe(III)], which acts as the final electron acceptor under environmental conditions (6, 7) in a process known as Feammox (8–10). Similarly to other metal reducing bacteria, *Acidimicrobiaceae* bacterium A6, a type of Actinobacteria, has the ability to use other sources of electron acceptors (11). The Actinobacteria phylum is commonly present in microbial community composition analysis of biomass associated to electrodes (12–17), but its role on the electrodes is rarely analyzed, most likely because it is not amongst the most abundant. To the best of our knowledge, to this date, there is only one report of an electrogenic Actinobacteria, genus *Dietzia*, a heterotrophic bacteria isolated from an intertidal zone at the Río de la Plata River (18). A6 is an iron reducer (7) that can use AQDS (anthraquinone-2,6-disulfonate), a humic acid analogue, as electron shuttle (10). Therefore, these A6’s characteristics opened up the possibility of it also being an ERB.

Lithoautotrophs are microorganisms that use inorganic compounds for their energy source and CO_2_ as their carbon source. This type of microorganisms are usually studied as part of the communities that develop at the cathode because they can be electrotrophs, i.e. they can uptake electrons directly from the cathode as their energy source, and they thrive on the CO_2_ formed by the oxidation of organic matter by the ERB (19, 20). The microbial communities that develop at the cathode are as highly diverse as the communities that develop at the anode, and much of the microbial groups found at the anode as ERB have also been found at the cathode (17, 21), some of them as proven electrotrophs, including *Geobacter* species (20, 22). Among the phyla present on both, anode and cathode, one can usually find Actinobacteria.

Microbial fuel cells (MFCs) and microbial electrolysis cells (MECs) are two reactor configurations that utilize electrogenetic microorganisms for renewable energy production, bioelectricity generation and pollutant degradation. In particular, MFCs coupled to constructed wetlands (CWs) have been used as devices to explore the possibility of treating wastewater and producing electricity simultaneously (23, 24). By incorporating MFCs in planted constructed wetlands, MFC operation can be promoted by the oxygen excreted by plant roots (25), resulting in stratified redox conditions that develop in wetland soils (26).

Given that the Feammox process has been found in multiple submerged sediments (7, 8, 27–29) and that A6 was isolated from wetland sediments (30), studying it in the field and in CW is advantageous to understand and characterize this bacterium. In this study, electrodes were installed in multiple locations of a forested riparian wetland as well as in laboratory CW mesocosm to investigate A6 as a potential ERB. The field locations provide insightful information regarding A6’s natural and electrode-enhanced thriving conditions, whereas the MFCs coupled CW mesocosms make controlled conditions available to better understand the field findings.

The objectives of this study were to (i) investigate if *Acidimicrobiaceae* bacterium A6 could colonize electrodes placed into sediments of a location where *Acidimicrobiaceae* bacterium A6 has been previously detected, and thus enhance its number with respect to the surrounding population, and (ii) to analyze its ability to transfer electrons to an electrode by determining current increments in CWs with embedded electrodes in response to bioaugmentation with *Acidimicrobiaceae* bacterium A6. These findings expand the knowledge of the diversity of ERB by including a member of the previously overlooked Actinobacteria, and allow for the possibility of practical applications for *Acidimicrobiaceae* bacterium A6 for the treatment of NH_4_^+^ contamination in anoxic systems using electrodes as stable, long-term terminal electron acceptors.

## RESULTS

### *Acidimicrobiaceae* bacterium A6 quantification

#### A6 quantification in the field study

Bacterial count by qPCR confirmed our initial hypothesis that the number of the A6’s population could be enhanced on the electrodes’ surface because the bacteria may have the ability to use electrodes as electron acceptors in the same manner that other FeRB do. The average count of A6 on the electrode’s biofilm was orders of magnitude higher than the average count of the bacterium in the soil (5.21E+7 copies of DNA/m^2^ on the electrode vs 2.33E+3 copies of DNA/m^2^ in the soil). Furthermore, in order to confirm that the electrodes were also being colonized by other electrogenic bacteria, we choose to quantify the genus *Geobacter*, which has become a model organism for the study of electrogenic bacteria. Both organisms, A6 and *Geobacter* spp., had significant higher populations on the biofilm formed on the electrodes surface than on the surrounding soil (Figure 1) (Welch t-test, p < 0.05 for A6 and p < 0.001 for *Geobacter*). On average, across all sites, the biofilm quantification for A6 and *Geobacter* resulted in ~4 orders of magnitude higher bacterial counts on the electrodes than on the surrounding soil (Table 1), which indicates that the number of A6 was clearly enhanced by the electrodes. However, we could not always see a trend in biomass being higher at the deeper electrodes than on the shallow electrodes as initially hypothesized. We had initially assumed that the surroundings of the shallow electrode would be the more oxidized, thus acting as the cathode, and the deeper the more reduced, hence acting as an anode. Nonetheless, because the electrodes were placed in the field for 52 days without any interference, during which 19 days of rain were recorded with a monthly average precipitation of 42 mm in June and 144 mm in July 2016 (Figure S1), the redox state of the soils could have shifted, thus inverting the polarity of the electrodes due to the fluctuation in the water table. Such conditions could have favored the colonization of electrode-reducing bacteria on both electrodes, the deep as well as on the shallow ones at different times. Furthermore, two current measurements were taken in the field between the deep and shallow electrodes of each set, on the first and final day. For set 1, current increased from 0.40 to 40.45 μAmps, set 2 from 0.10 to 0.40 μAmps, set 3 from 0 to 2 μAmps, and set 4 from 0.50 to 0.75 μAmps. For Site 5-6, current measured on set 5 decreased from 7 to 0.66 pAmps, and set 6 inverted its current from 8 to -0.50 μAmps, thus indicating that electrode polarity could have inverted during the time when the electrodes were in the field. Therefore, the redox potential profile in each of the CWs was continuously monitored to stablish which electrodes were working as the cathodes and as the anodes.

**Figure 1.**
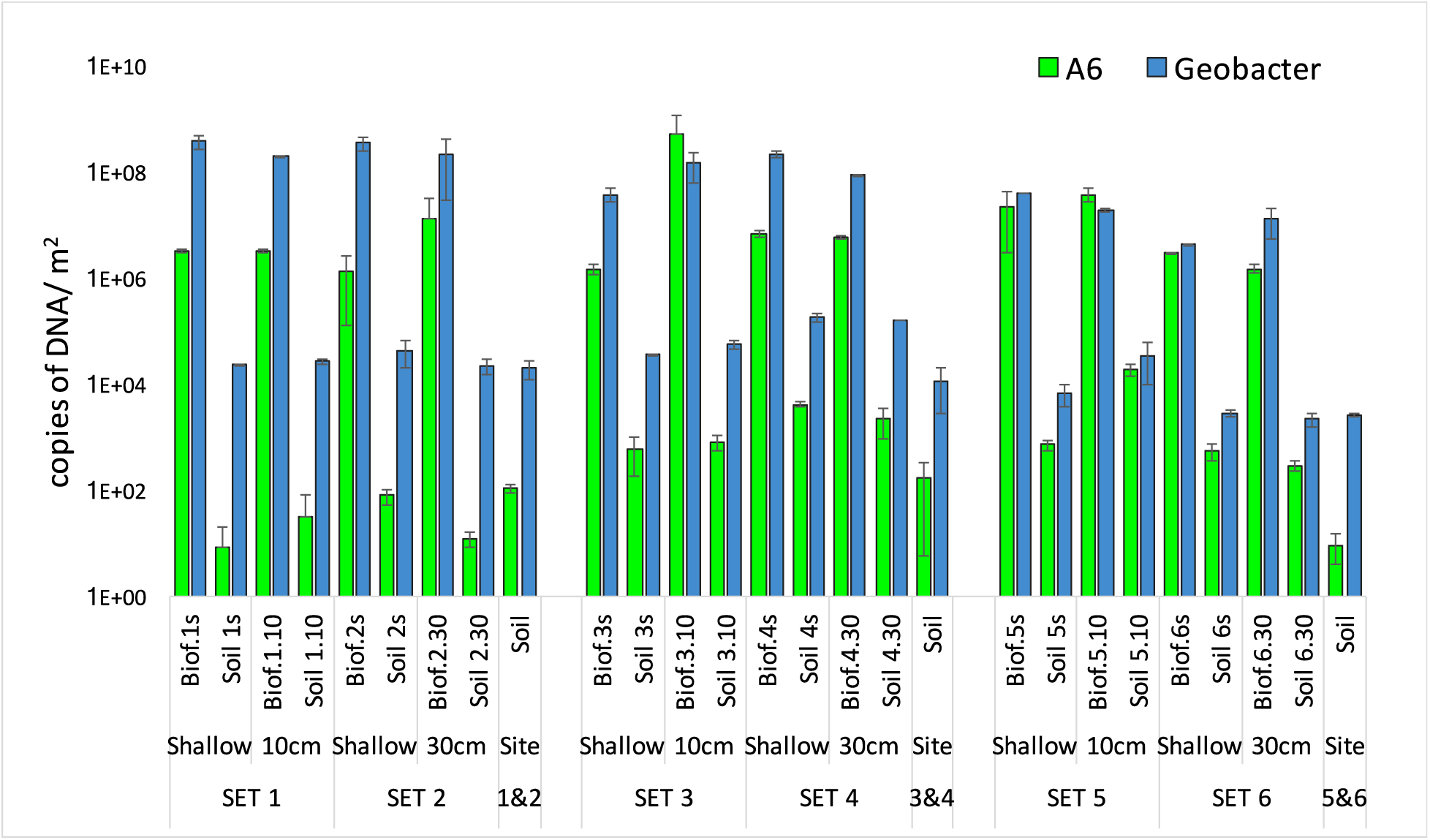
*Acidimicrobiaceae* bacterium A6 and *Geobacter* spp. quantification from biofilm formed on the electrodes and soil samples taken from 3 field electrode sets.

**Table 1.**
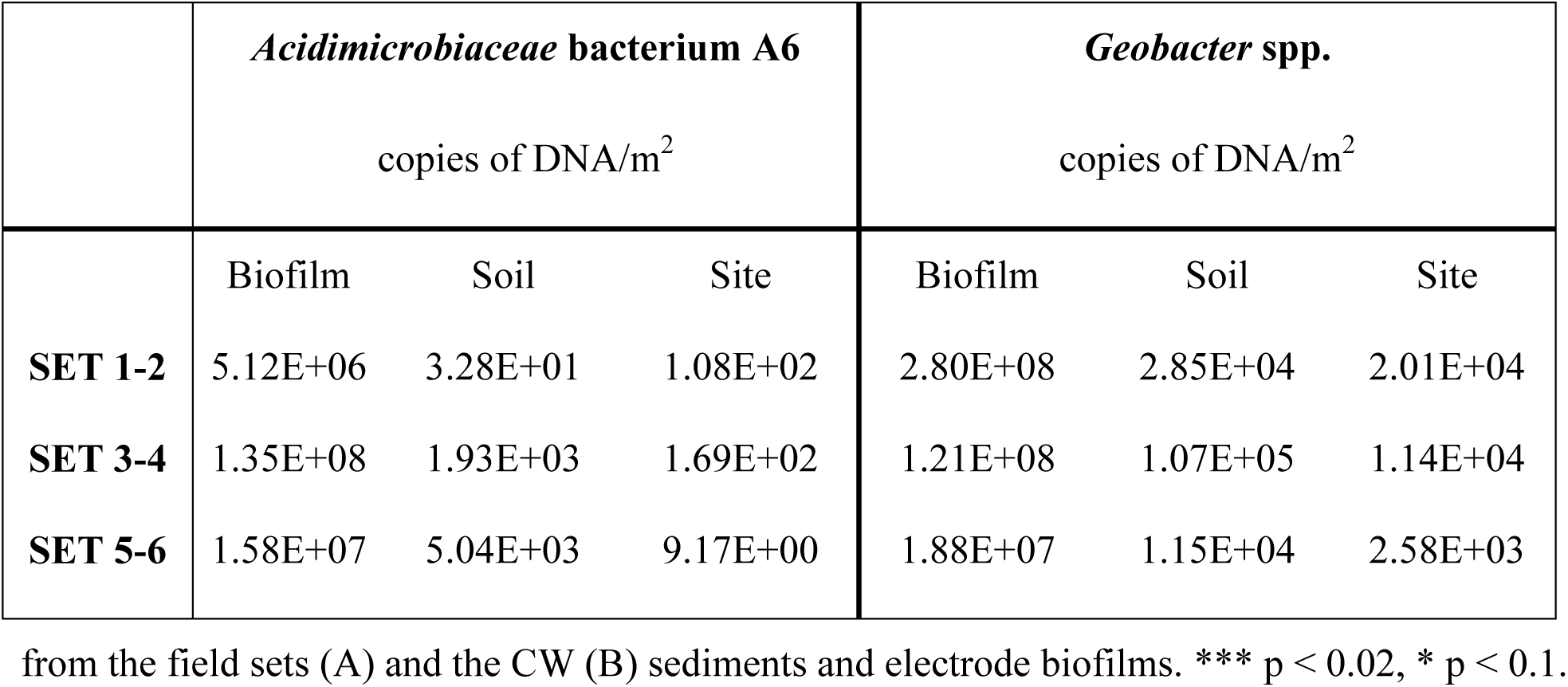
Average number of copies of DNA /m^2^ quantified for *Acidimicrobiaceae* bacterium A6 and *Geobacter* spp. from samples obtained from biofilm, soil surrounding the electrodes, and soil samples for each site.

|  | *Acidimicrobiaceae* bacterium A6 |  |  | *Geobacter* spp. |  |  |
| --- | --- | --- | --- | --- | --- | --- |
|  | copies of DNA/m<sup>2</sup> |  |  | copies of DNA/m<sup>2</sup> |  |  |
|  | Biofilm | Soil | Site | Biofilm | Soil | Site |
| **SET 1-2** | 5.12E+06 | 3.28E+01 | 1.08E+02 | 2.80E+08 | 2.85E+04 | 2.01E+04 |
| **SET 3-4** | 1.35E+08 | 1.93E+03 | 1.69E+02 | 1.21E+08 | 1.07E+05 | 1.14E+04 |
| **SET 5-6** | 1.58E+07 | 5.04E+03 | 9.17E+00 | 1.88E+07 | 1.15E+04 | 2.58E+03 |

from the field sets (A) and the CW (B) sediments and electrode biofilms. \*\*\* p < 0.02, \* p < 0.1.

#### A6 quantification in the CW mesocosm study

The population of A6 (copies of DNA / m^2^) is shown in Figure 2 for all samples from both CW mesocosms. The A6 population is higher in the high Fe mesocosm than in the low Fe mesocosm either on the CW sediments or on electrode biofilms (p < 0.05). Furthermore, the A6 population on the electrode biofilms is always 2 – 3 orders of magnitude higher than in their surrounding sediments for the deeper electrodes (depth > 15 cm) in both mesocosms.

**Figure 2.**
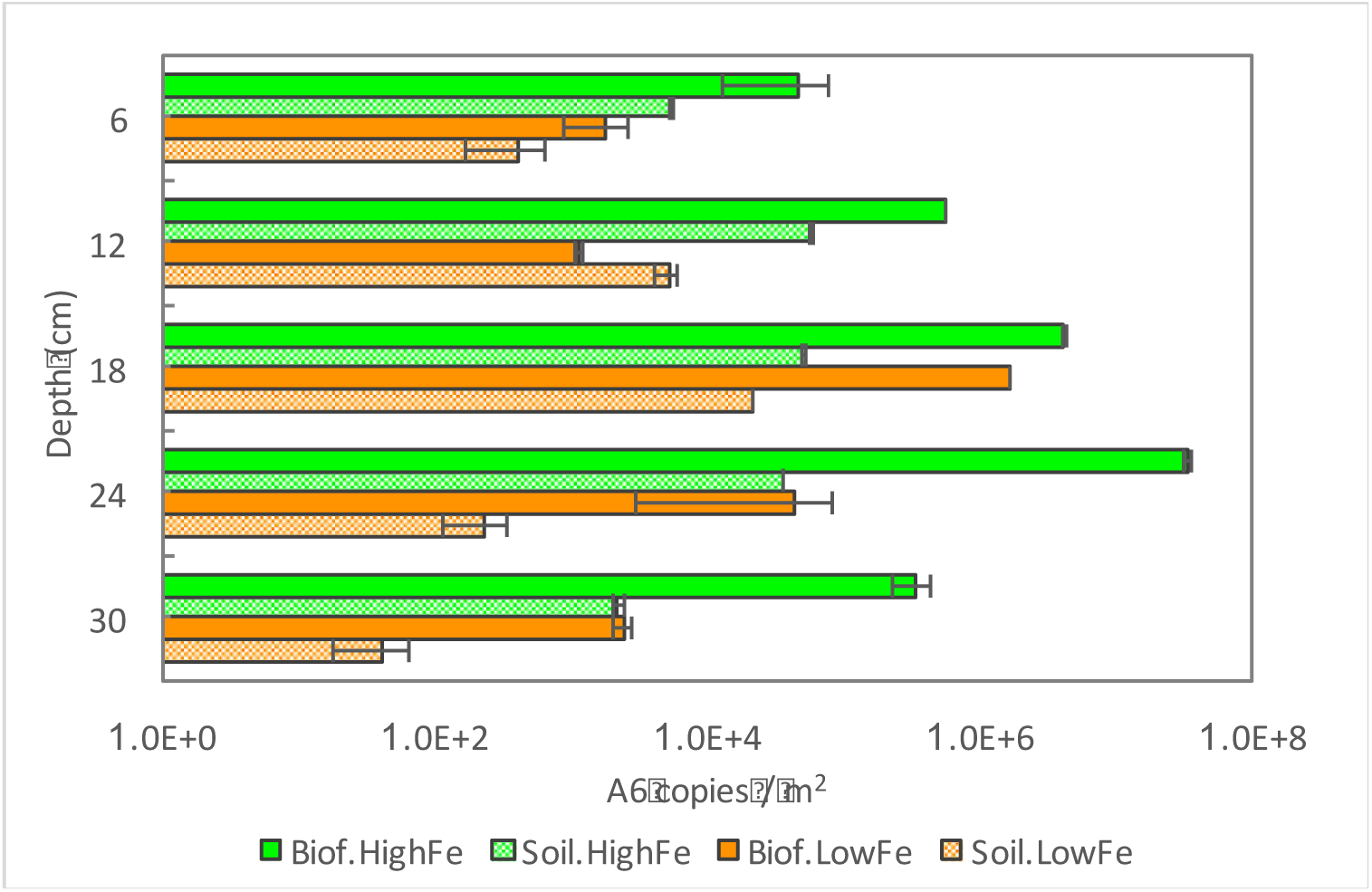
*Acidimicrobiaceae* bacterium A6 populations in the CW sediments or the electrodes in CW mesocosms. Error bars are upper and lower values of qPCR measurements.

Electrode 1 (at depth = 6 cm) was designated as cathode at the beginning, but as the redox potential profile in the CW developed, the direction of current between the electrode 1 and electrode 2 (at depth = 12 cm) reversed around day 35. Hence, on day 59 electrode 2 was designated as cathode and the wires were reconnected to form electrode pairs between electrode 2 and other electrodes. Figure S2 shows that the ORP at the location of electrode 2 was the highest before the injection of A6 enrichment culture. As the electrodes below 15 cm (electrodes 3 – 5) had lower oxidation-reduction potential through the entire experiment, those electrodes always operated as the anodes. The tendency of A6 colonization was observed on the anodes (for A6 count on electrode vs. in soil at depth >15 cm, p < 0.05) whereas the two shallower electrodes, which each operated as cathodes for a certain period, did not always show larger numbers of A6 on these electrodes compared to the surrounding soil, and no statistically significant difference in the count number between those samples was found (for A6 count on electrode vs. in soil at depth < 15 cm, p > 0.05). These results confirm the higher affinity of A6 for the electrode (anode) that remained in the more reduced soil throughout the experiment over the electrode (cathode) in the more oxidized soil. The well-controlled and monitored CW mesocosms were able to provide insights that were not clearly resolved by the field studies.

### Phylogenetic analyses and microbial community structure

The phylogenetic diversity at the phylum level found in the biofilm and soil samples taken from the field as well as the CW mesocosms studies (Figure 3) show that Proteobacteria and Acidobacteria represent on average more than 70% of the diversity found in all samples, followed in abundance by Chloroflexi and Bacteroidetes in the field study, and Firmicutes and Bacteroidetes in the CW mesocosm study. These highly abundant groups make up more than 80% of the population found in all samples. All these phyla are commonly found in soil and in bioelectrochemical systems due to their ERB ability (12, 13, 31, 32), therefore, they are common subjects of study. Actinobacteria, the phylum to which *Acidimicrobiaceae* bacterium A6 belongs to, represents as little as 2, 4, 5 and 3.5 % of the relative abundance found at the three different sites and CW mesocosms respectively. Nonetheless, Actinobacteria ranks in the top 5 most abundant phyla found in each field site and the CW mesocosms, and makes it to the third position for some electrode biofilm samples from the field study.

**Figure 3.**
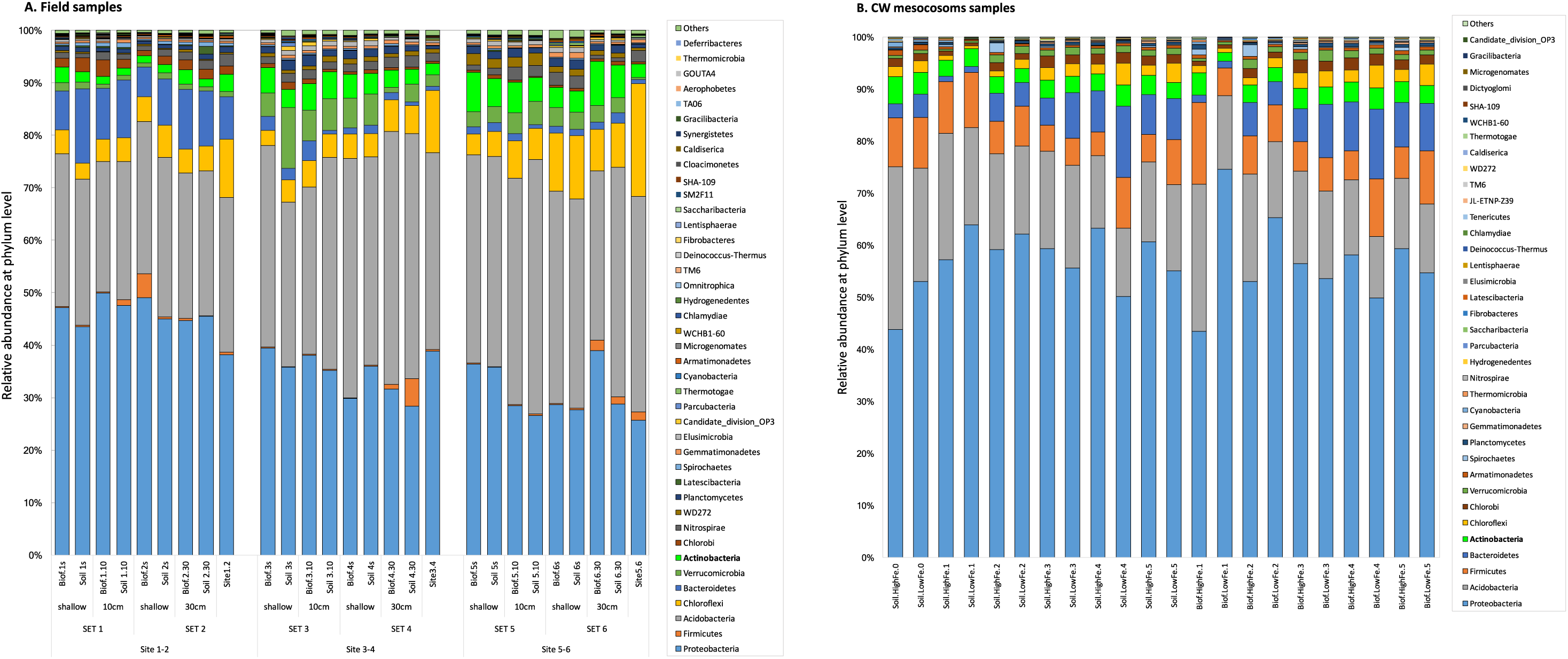
Microbial community composition at the phylum level of biofilm and soil samples from electrode pairs from 3 different field locations (A) and CW mesocosms (B). Actinobacteria phylum, highlighted in yellow, to which *Acidimicrobiaceae* bacterium A6 belongs, is amongst the most abundant phyla in all samples.

#### Field experiment phylogenetic analyses

The Actinobacteria phylum contains *Acidimicrobiaceae* bacterium A6, described as an unidentified_*Acidimicrobiales* at the genus level in the samples from the field sites, because its 16s rDNA sequence was not available in the public data bases at the time of the field study. The OTU annotated as unidentified_*Acidomicrobiales* had ≥ 97 % sequence identity with A6, thus confirming the presence of this Feammox bacterium in our samples. A total of 316 genera were annotated in the phylogenetic analysis, however, between 51% (site 1-2) to as much as 69% (site 5-6) of the OTUs could not be classified at this level, thus, they were added to the “others” category. Among the top 100 most abundant genera, the unidentified *Acidomicrobiales* ranked 56^th^ (Figure 4). The genera with the highest relative abundance at site 1-2, characterized by its waterlogged condition, were *Sideroxydans* (Proteobacteria), an Fe(II) oxidizer (33), and *Geothrix* (Acidobacteria), a known ERB (34). At sites 3-4 and 5-6, the most relative abundant genera were the *Bryobacter* (Acidobacteria) an aerobic heterotroph, candidatus_*Solibacter* (Acidobacteria), *Acidibacter* (Proteobacteria) an FeRB, the autotroph *Acidothermus* (Actinobacteria), and *Sorangium* (Proteobacteria). Other Fe cycling bacteria found among the top 100 most abundant genera are *Acidiferrobacter*, *Anaeromyxobacter*, *Ferritrophicum*, *Geobacter*, *Gallionella*, *Desulfobulbus*, and *Georgfuchsia*, all from the Proteobacteria phylum.

**Figure 4.**
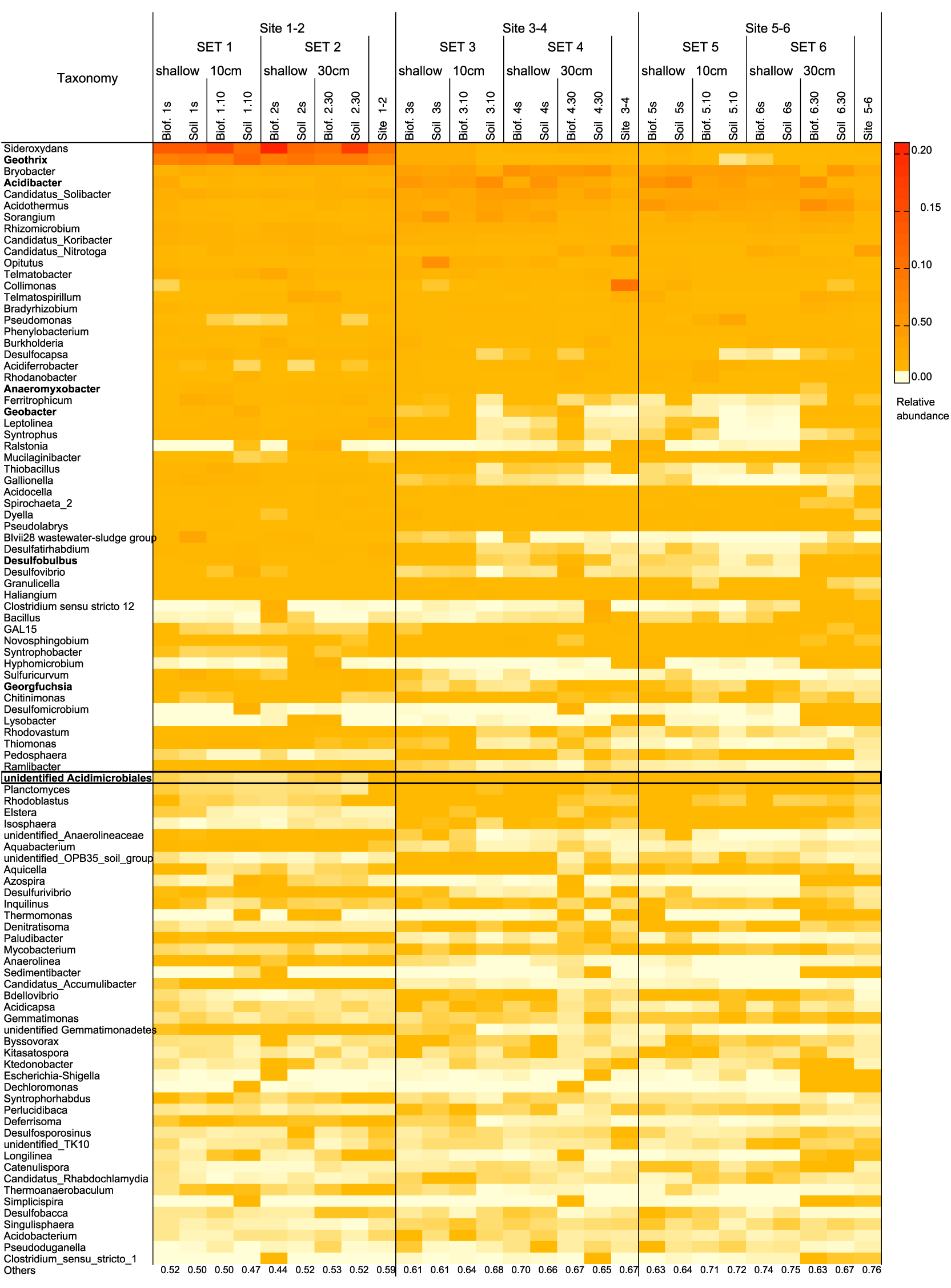
Relative abundance of the top 100 most abundant genera in biofilm and soil samples from the electrode sets placed in the field. *Acidimicrobiaceae* bacterium A6 had 97% identity with the unidentified_*Acidomicrobiales* which ranked 56^th^ in abundance. In bold other Fe-reducing bacteria.

#### CW mesocosm phylogenetic analyses

A6 was identified in the CW sediments and electrode biofilms, and A6 ranked 89th on average for all (soil and electrode biofilm) CW mesocosm samples among the top 300 genera. Unclassified OTUs comprised 26.3 – 51.6 % of the total sequences for CW soil samples and 17.4 – 49.5 % of the total sequences for electrode biofilm samples. The two genera with the highest relative abundance in the CW mesocosms for all (soil and electrode biofilm) samples were *Thiomonas* and *Burkholderia*, both are Proteobacteria. In the CW soil samples the third most abundant genus was *Telmatobacter* (Acidobacteria), a group of anaerobes and chemo-organotrophs, while in the CW electrode biofilm samples the third most abundant genus was *Geobacter* (Proteobacteria).

The relative abundance of A6 in all CW samples is about 0.1 – 12.8 % of *Geobacter spp*., which can be enriched on the anodes (3). Although A6 are not as abundant as *Geobacter spp*. (Figure S3) in the CW mesocosms, their population was still enriched on the anodes compared to the surrounding sediments (Figure 2). In addition to *Geobacter*, 10 other genera that include known electrogenic bacteria species were also detected in all CW mesocosm samples among the top 100 genera, including *Geothrix*, *Desulfobulbus*, *Desulfovibrio*, *Pseudomonas*, *Clostridium* (2), *Desulfotomaculum* (35), *Enterobacter* (36), *Bacillus, Rhizomicrobium* (37) and *Anaeromyxobacter* (31). Most of the mentioned electrogenic bacteria are more abundant in the electrode’s biofilms than in the nearby soil samples (paired two sample t-test, p < 0.05; Table S3). In fact, the electrogenic bacteria form a substantial portion of the microbial community from the CW mesocosm samples, making up 3.0 – 8.5 % of the total sequences for the CW soil samples and 4.5 – 14.4 % for CW electrode biofilm samples. Those electrogenic bacteria that are enriched on electrodes compared to the nearby soil are all much more abundant than A6, resulting in lower relative abundance of A6 on the electrode biofilms compared to the nearby soil, even though the numbers of A6 are higher on the electrodes than the soil.

### Current pulse after the injection of A6 enrichment culture into the CWs

Though the high and low Fe level CW mesocosms had similar current profiles before the injection of the A6 enrichment culture, the current profiles right after the injection showed a noticeable difference (Figure 5). The electrical current between electrode pairs in the low Fe CW mesocosms increased after five days following the A6 enrichment culture injection, and then descended to the previous level after 50 days. However, currents between electrode pairs in the high Fe CW mesocosms remained within a similar range as prior to the A6 enrichment culture injection, and then decreased at around the same time when the pulse in current in the low Fe CW mesocosm disappeared. The different responses indicate that more electrons were transferred through the electrode pairs in the low Fe mesocosm than in high Fe mesocosm as the same amount of bacterium A6 was introduced into both mesocosms.

It should be noted that the samples for DNA extraction were taken almost four months after the injection of A6 and three months after the current pulse disappeared in the low Fe mesocosm. Therefore, phylogenetic results and A6 numbers discussed above may not properly capture the microbial community at the time of the pulse in the current.

**Figure 5.**
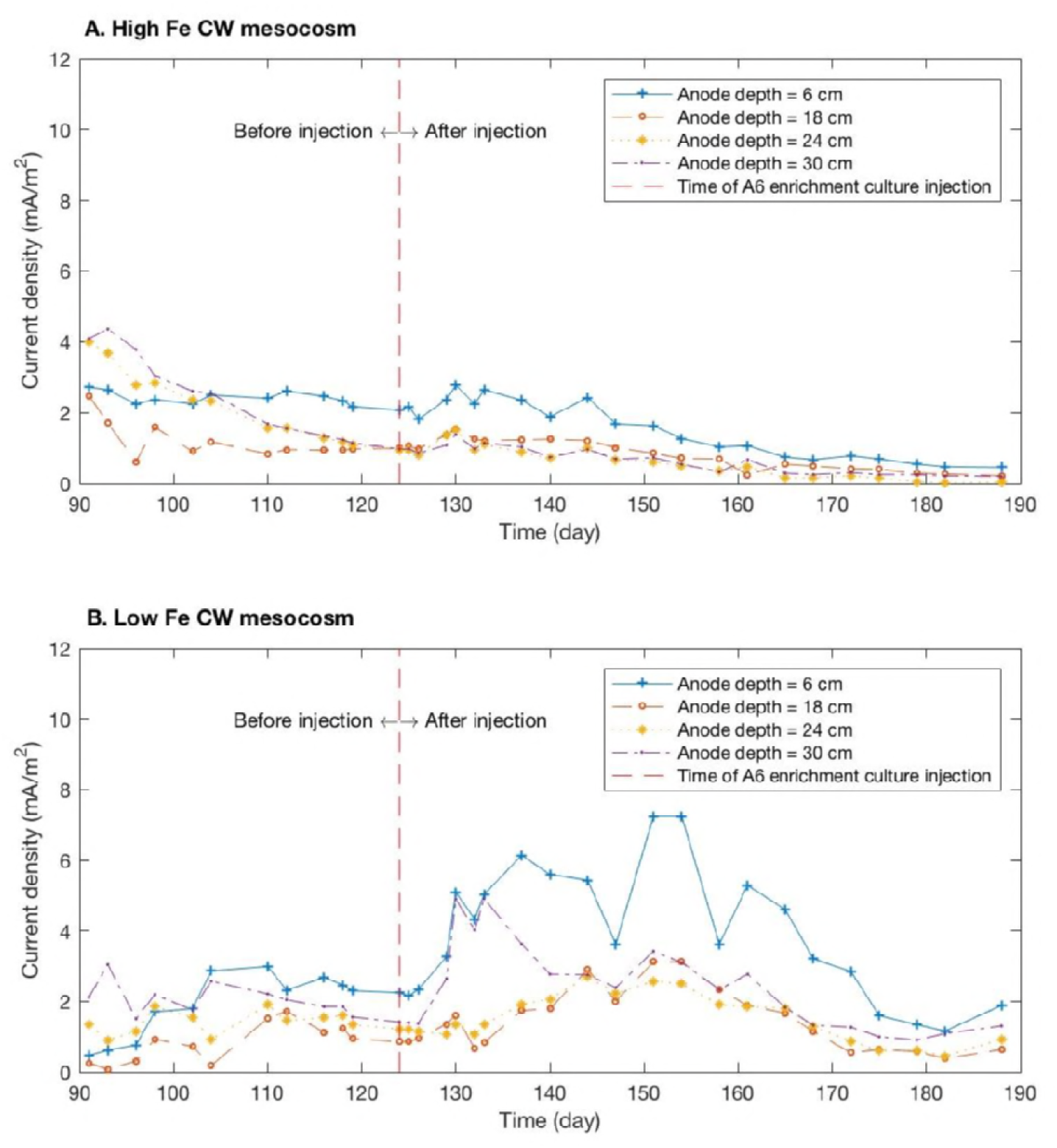
Current density profiles for high Fe CW mesocosm (A) and low Fe CW mesocosm (B) one month before and three months after the injection of the A6 enrichment culture. Because of the ORP development, the second electrodes (at depth 12 cm) had the highest redox potential when A6 enrichment culture was injected. Therefore, the electrodes at depth 12 cm in both CW mesocosms were connected as cathodes. The currents were measured for electrode pairs with different anode depths.

## DISCUSSION

### A6 colonization and electron transfer to electrodes

The number of A6 quantified on the biofilm formed on the electrodes confirmed the hypothesis that this bacterium is able to colonize electrodes to use them as an alternative electron acceptor to Fe(III), thus enhancing its number compared to its surroundings (Figure 1). The CW permitted us to stablish A6’s preference over the electrodes in the more reduced soil (anodes) than over the rest of the environment (Figure 2). The consistent trend of A6 enrichment on electrodes was found on the anodes compared to surrounding sediments but not always on the electrodes that have been operated for a time period as cathodes (electrode 1 and 2 in the CW). This indicates that A6 is able to colonize and be enriched on the surface of anodes.

It is interesting to note that injection of A6 into the low Fe CW resulted in a pulse in current while this was not observed in the high Fe CW. This indicates that when little bioavailabe Fe was present, A6 showed a more immediate affinity to colonize and transfer electrons to the electrodes. The decrease in the current after about 50 days of the injection indicates that A6 numbers or activity on the electrodes in the low Fe CW decreased over time after the injection.

### *Acidimicrobiaceae* bacterium A6 and other Fe-cycling bacteria

A6 ranked 56^th^ in the relative abundance at the genera level in the field study samples, and much lower in the CW samples, being outranked by other FeRB, with most of which A6 showed negative correlations between their relative abundances in the field (Figure 6). This is not the case for the relative abundance with other non-metal reducing bacteria, also from field soil and electrode samples with which showed positive or no correlation (r = ~ 0.0) (Figure S4). When only the biofilms samples’ relative abundance correlations are analyzed, the correlation between A6 and *Collimonas* shifts from slight negative (r = -0.02) to a positive correlation (r = 0.45) (p > 0.1). For all the other genera, the trends of their correlations are maintained when all the data (biofilm and soil samples) is either pooled for analysis or separated by biofilm or soil samples only. This indicates, that in soils and electrode biofilms, A6 presence is negatively affected by most other Fe-cycling bacteria found in our samples. Whereas when the relative abundance of *Geobacter* is correlated with the other Fe-cycling bacteria, it shows a positive correlation with all except *Acidibacter* and *Acidiferrobacter*. These findings open up the need for further research to understand what drives these correlations and if they may indicate a competition for the electron acceptor between A6 and other FeRB.

**Figure 6.**
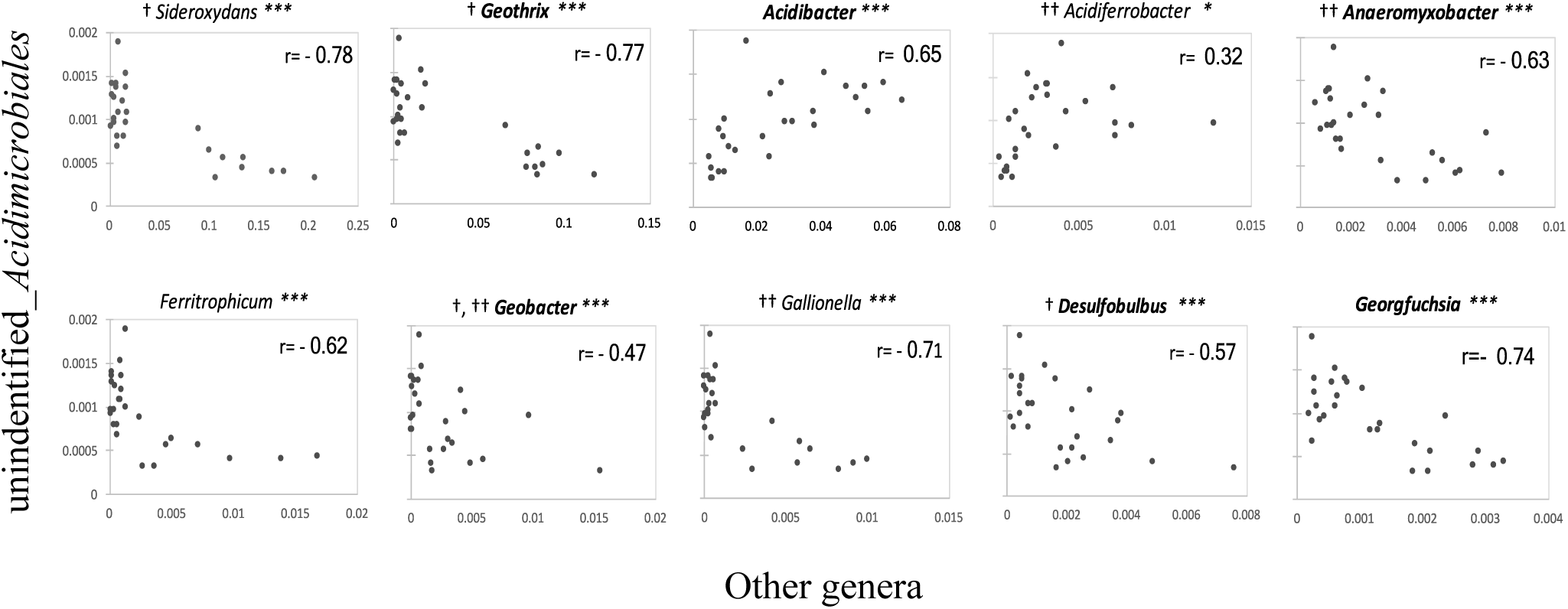
Correlation of the relative abundance between *Acidimicrobiaceae* bacterium A6 (unindentified_*Adicimicrobiales*) and Fe-cycling bacteria in biofilm and soil samples (n=27). Fe-oxidizing bacteria are in italics and Fe-reducing bacteria in bold italics. † Anode colonizer (22, 31, 34, 51), †† Cathode colonizer (31). ^∗∗∗^ p < 0.01, ^∗^ p < 0.1.

The higher relative abundance of A6 in the high Fe compared to the low Fe mesocosm (Figure S3) indicates that A6’s relative abundance positively responds to the increased Fe(III) level in the sediment. This positive correlation between A6 and Fe(III) has been previously reported in environmental samples (30). However, A6 and *Geobacter spp*. relative abundances are negatively correlated, with a correlation coefficient of -0.47 (p < 0.02) in field samples and - 0.38 (p < 0.1) in the CW (Figure 7), and this holds for both electrode biofilm samples and soil samples. This negative correlation should be taken into account when implementing A6 in bioelectrochemical system such as MFC, particularly those that feed on organic carbon as the electrode donors since these are systems where bacteria such as *Geobacter spp*. thrive and could affect A6’s population negatively.

**Figure 7.**
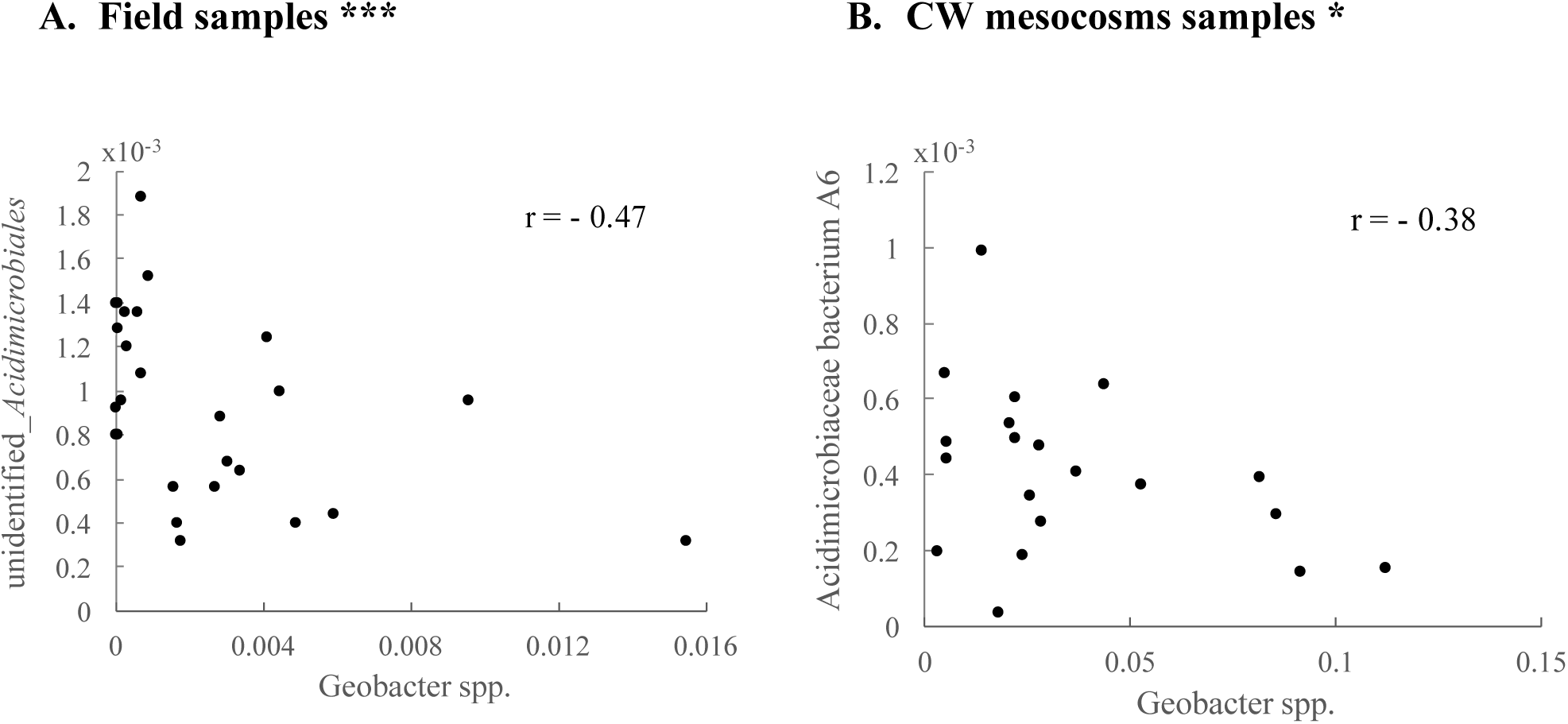
Correlation between Feammox bacteria and *Geobacter* spp. relative abundance. Data from the field sets (A) and the CW (B) sediments and electrode biofilms. ^∗∗∗^ p < 0.02, ^∗^ p < 0.1.

The results from this study show that *Acidimicrobiaceae* bacterium A6’s, which is an iron reducer, is capable of colonizing electrodes in the field as well as in constructed wetland mesocosoms, resulting for the conditions studied, in higher cell counts on the electrodes than on the soil. Thus, *Acidimicrobiaceae* bacterium A6 is a novel anaerobic litoautothroph from the Actinobacteria phyla capable of using electrodes as its terminal electron acceptor. Altogether, this work expands the knowledge of the diversity of electrogenic microorganisms beyond the commonly studied groups and opens up the possibility for applications of this bacteria in MFCs and MECs systems. However, further research is needed to elucidate what drives the different interactions between A6 and other FeER and ERB in order to optimize its applications in bioelectrochemical systems.

## MATERIALS AND METHODS

### Field electrodes construction and setup

Electrodes consisted of graphite plates [7.5 (L) × 2.5 (W) × 0.32 cm (H)], with a surface area per face of 18.75 cm^2^ (Grade GM-10; GraphiteStore.com Inc.). The plates were polished using sandpaper (grit type 400), sonicated to remove debris, cleaned by soaking in 1 N HCl overnight and rinsed three times in distilled water (38). Each electrode set was connected by a titanium (Ti) wire cleaned with sandpaper (ultra-corrosion-resistant Ti wire, 0.08cm in diameter, McMaster-Carr code 90455k32) by inserting the wire through two holes of size 0.08 cm drilled in each graphite plate to ensure a tight connection between the wire and the graphite plates to allow for low contact resistance <0.5 Ω. The Ti wire was long enough to allow for 10 or 30 cm separation between the graphite plates (Figure 8).

**Figure 8.**
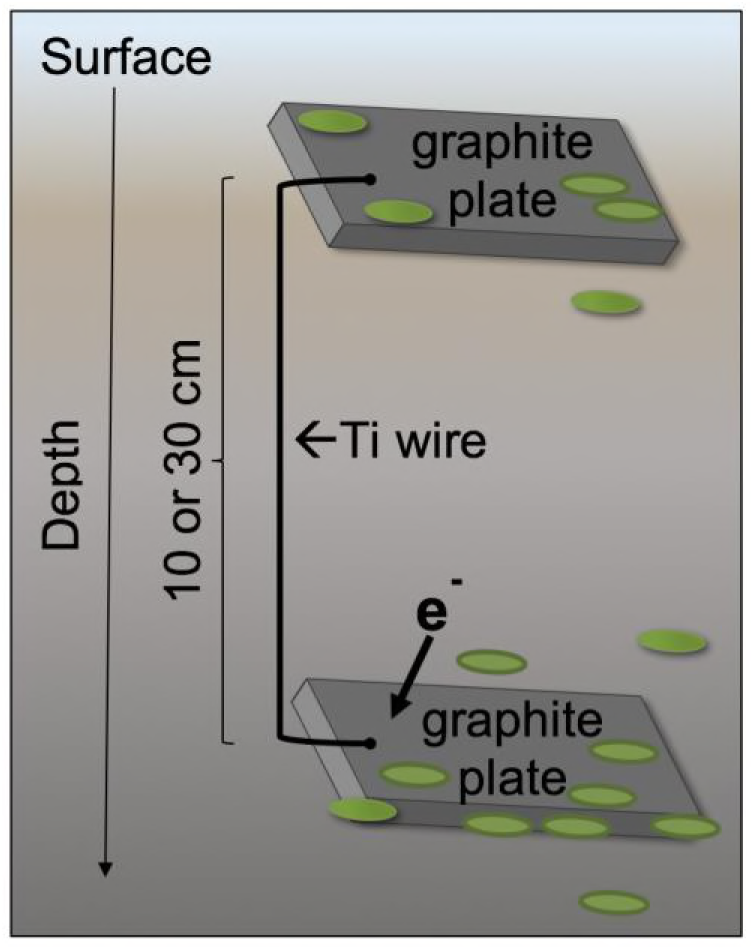
Schematic of an electrodes pair.

Two pairs of electrodes were placed at three different sites. Each pair consisted of a shallow electrode placed no deeper than 5cm into the soil, connected to another electrode with either 10 cm or 30 cm separation, i.e. a total of 6 sets (Table 2) in a temperate forested riparian wetland located at Assunpink Wildlife Management Area in New Jersey, USA. This is the location were the Feammox reaction was first discovered (8), and later the Feammox bacteria A6, was identified in samples from this site (7) and isolated (10). Detailed physicochemical characteristic of the soil have been described in previous studies (8, 39). Electrodes sets 1 and 2 were place in a fully flooded location, sets 3 to 6 were placed in a wet but unsaturated location. The electrode sets were left in the field for 52 days between June 13 and August 03, 2016.

**Table 2.**
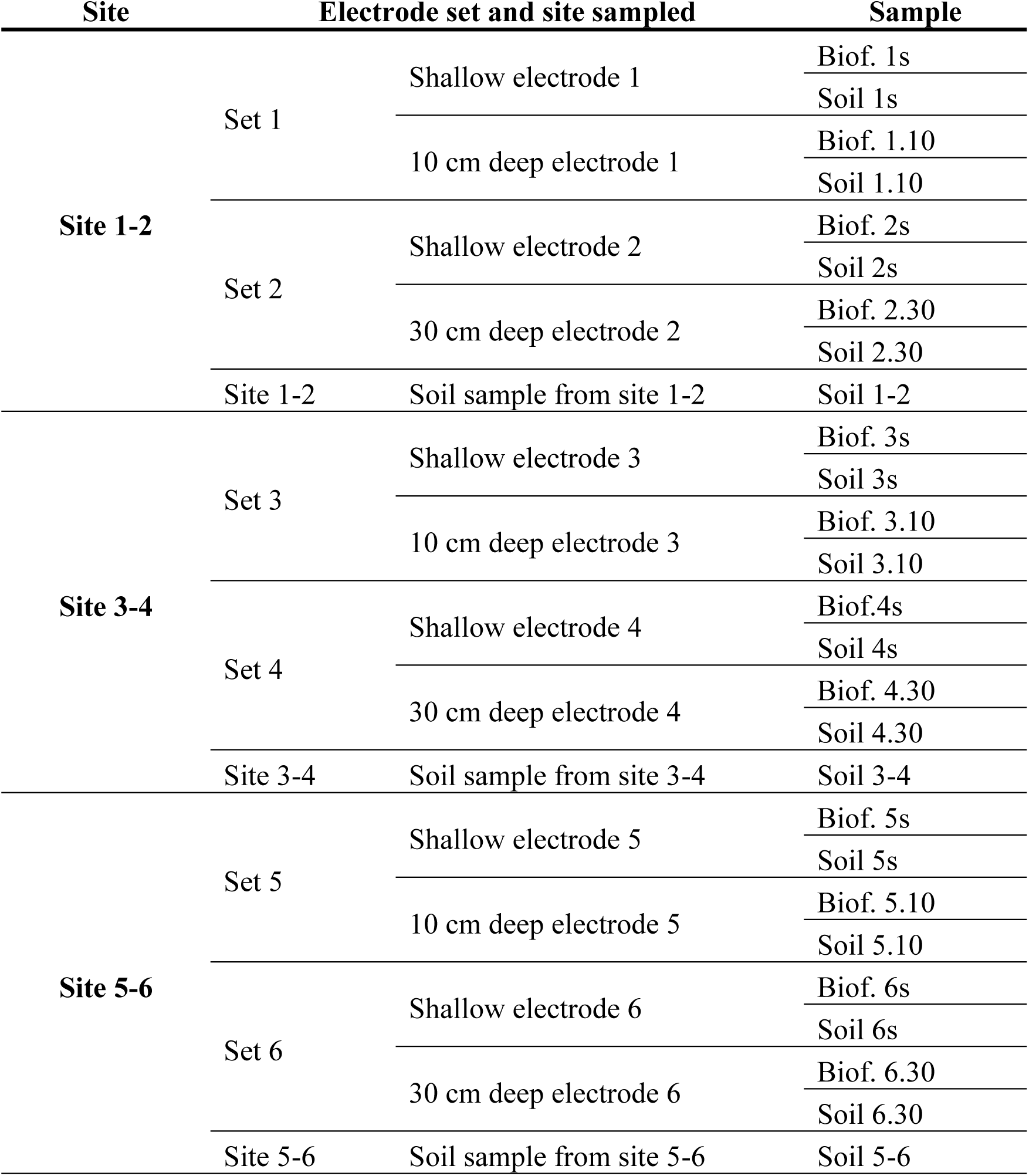
Description of biofilm (Biof., sample #, depth) and soil (Soil, sample #, depth) samples taken from the electrode pairs located at each location.

### Field electrodes recovery and sampling

After 52 days, the electrode pairs were recovered by digging them out of the soil and placing each electrode individually in a sealed bag. All the electrodes were surrounded by soil. Furthermore, a soil sample was taken from a depth of ~20 cm from each site and placed in a sealed bag. All samples were transported to the laboratory within 2 hours and immediately stored at 4°C until processed for analysis.

The samples obtained from the electrodes and soil are enumerated in Table 2. To analyze the biomass attached to the electrodes, first, the loosely bound soil was removed by gently shaking the electrode. Second, duplicate samples were taken from the soil layer (< 2 mm thick) still surrounding the electrode. Third, duplicate samples of the biomass formed on the electrodes’ surface together with some graphite were removed using a sterile cutting blade (see details in Supplemental Materials, Table S1. A). DNA was extracted from all samples and then used for determining, quantifying, and comparing their microbial composition.

### Constructed wetland mesocosms and electrodes set up

Controlled conditions were required to gain further insights into the electrogenesis of A6. Therefore, electrode pairs were installed in two constructed wetland (CW) mesocosms that were operated in a growth chamber (Environmental Growth Chambers, www.egc.com). The constructed wetlands were designed as continuous up-flow mesocosms with water surface above the sediment. Standard-wall PVC pipes, pipe fittings (pipe size 6, inner diameter 6 inch, McMaster-Carr code 48925K25, 4880K852 and 4880K131) were used to assemble the mesocosms. The dimension of the mesocosms and the design of sampling ports are shown in Figure 9. The inflow was injected from the bottom and an opening with 1-inch diameter was drilled for the effluent to maintain constant water levels. Along the longitudinal axis were five sampling ports, spaced 6 cm apart from each other, and lysimeters were used for pore water sampling from the CW mesocosms. Nylon meshes (70 μm opening, opening area 33%) were placed at 15 cm and 40 cm depth to separate the root zone/non-root zone respectively. At the bottom of the mesocosms, glass beads were used as bed material to disperse the inflow evenly. The CW substrate was a mixture of ASTM standard 20-30 sand, peat moss, and wetland sediments from Assunpink Wildlife Management Area in New Jersey, USA (the same location where the field electrodes were placed) with 1: 1: 0.5 ratio (by weight).

To investigate if A6 could colonize the electrodes and transfer electrons onto anodes, five graphite electrodes were installed in each mesocosm at the same depths as the five sampling ports. The electrodes were made of rectangular graphite rods [10.16 (L) × 1.28 (W) × 1.28 cm (H)]. The preparation of the electrodes and wires was the same as described above. Each of the four electrodes that were placed into the more reduced zones (anode) was then connected via a cleaned titanium wire to the electrode that was in the most oxidized zone (cathode) of the CW as shown in Figure 9.

Synthesized Fe_2_O_3_0.5H_2_O (2-line ferrihydrite) was added to one of the CW mesocosms to elevate its initial Fe(III) level. A concentration of 500 mg Fe(III) / kg moist sediment was added to the high Fe CW mesocosm and mixed thoroughly with the sediments, whereas no extra Fe(III) source was added to the low Fe CW mesocosm sediment. The method for synthesizing 2-line ferrihydrite was modified from Schwertmann and Cornell (2000) (40). After blending substrate thoroughly, Fe(III) was measured for high Fe CW mesocosm [~ 2.9 g Fe(III) / kg dry soil] and low Fe CW mesocosm [~ 1.7 g Fe(III) / kg dry soil] substrates. Since the wetland sediments and peat moss both had some Fe, even the CW that was not augmented with ferrihydrite had Fe. As ferrihydrite is amorphous and highly bioavailable, the Fe(III) source for A6 would mostly come from the ferrihydrite added in the CW substrate. Examination of several Fe(III) sources as the electron acceptor for A6 showed that this ferrihydrite yielded the highest Feammox activity among the Fe(III) sources studied (10).

To inoculate the CW substrate with A6, 250 mL of an A6 enrichment culture (10^9^ – 10^10^ CFU/g sludge, 70% A6 in biomass) was added to the CW substrate for each column. The substrate was then thoroughly mixed prior to loading into the mesocosm columns.

Each mesocosm was planted with four *Scirpus actus* plants (bulrush), obtained from Pinelands Nursery and Supplies, Columbus, NJ, USA.

### CW mesocosm operation

A half-strength modified Hoagland nutrient solution (41) with a high NH_4_^+^ concentration (100 mg/L NH_4_^+^-N) was pumped into the mesocosms at a flow rate of ~1.5 L/day. Before pumping into the CW mesocosms, the nutrient solution was mixed with 1 M acetic acid to increase the dissolved organic carbon and aid in the removal of the dissolved oxygen as well as lower the pH of the nutrient solution as the Feammox process requires acidic conditions (30). Each liter of half-strength modified Hoagland nutrient solution contained 6.64 mL 1 M NH_4_Cl, 0.5 mL 1 M NH_4_H_2_PO_4_, 3.0 mL 0.5 M K_2_SO_4_, 2.0 mL 1 M CaCl_2_, 1.0 mL 1M MgSO_4_, 0.5 mL micronutrient stock solution and 0.125 mL iron stock solution. The micronutrient stock was made by dissolving 2.86 g H_3_BO_3_, 1.81 g MnCl_2_4H_2_O, 0.22 g ZnSO_4_ 7H_2_O, 0.08 g CuSO_4_5H_2_O and 0.02 g H_2_MoO_4_H_2_O in 1 L of deionized water. The iron stock solution was made by adding 500 mL 49.8 g/L FeSO_4_7H_2_O solution slowly to the potassium EDTA solution (26.1 g EDTA in 286 mL water with 19 g KOH), aerating the mixture overnight while stirring and making the final volume 1 L.

The mesocosms and pumps were placed in a growth chamber that is simulating the summer climate of New Jersey (Table S2). Since the A6 enrichment culture was blended in open-air with the CW substrate and the CW mesocosm conditions were rather oxidized at the beginning of their operation, it was uncertain that a viable A6 population did get established in the mesocosms. Therefore, after four months of operation, when the mesocosm became more reduced, another A6 enrichment culture (~ 10^7^ A6 count/mL culture, 250 mL per CW mesocosm) was injected from the bottom into each CW on day 124 to ascertain the colonization of A6 in the mesocosms. This procedure also allowed to determine if a spike in A6 numbers would result in an increase in electrical current.

### CW mesocosm sampling and dismantlement

Oxidation-reduction potential (ORP) in the CWs was measured by taking water samples from sampling ports and collecting effluents from the top of the CWs. At the end of the experiment, the CW mesocosms were dismantled and soil samples were taken for analysis of microbial communities. Soil samples were collected at depths bracketing the sampling ports (6 – 9 cm, 12 – 15 cm, 18 – 21 cm, 24 – 27 cm, 30 – 33 cm) as well as at the top layer of soil (0 – 3 cm). For each depth, a 1 – 1.5 g wet soil sample was collected and frozen at -20 °C before proceeding with the DNA extraction. Electrodes installed in the CW mesocosms were removed carefully and biofilm samples from the electrodes were obtained using the same procedure as described above (see Supplemental Materials, Table S1. B for details). Since soil and electrode sampling requires sacrificing the mesocosms, no soil samples were collected during the operation of the CW mesocosms. Samples obtained from the CW mesocosm sediments and electrodes are enumerated in Table 3.

**Table 3.**
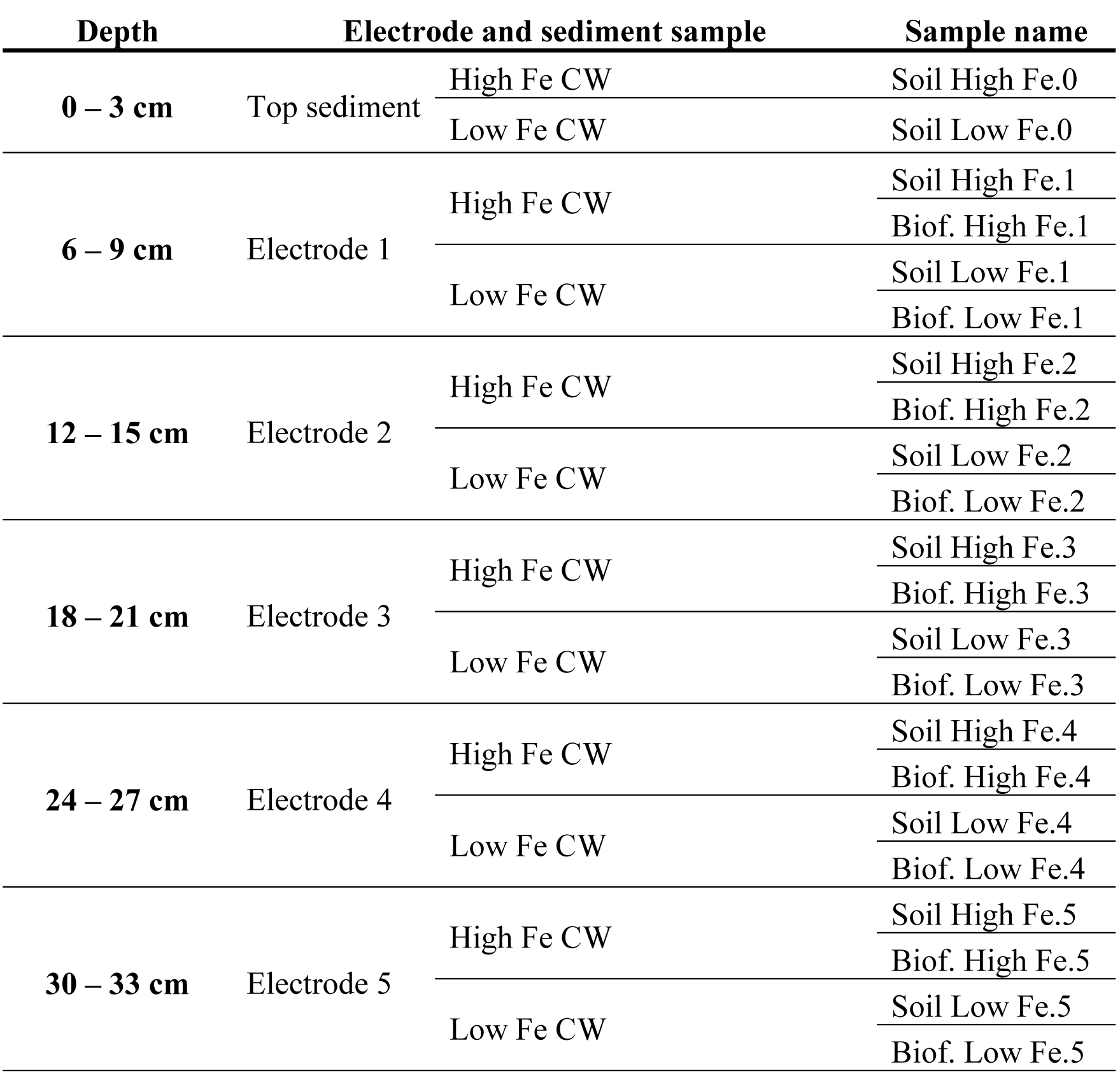
Description of biofilm (Biof., Fe treatment, electrode #) and soil (Soil, Fe treatment, electrode #) samples taken from the CW mesocosms.

| Depth | Electrode and sediment sample |  | Sample name |
| --- | --- | --- | --- |
| 0 – 3 cm | Top sediment | High Fe CW | Soil High Fe.0 |
|  |  | Low Fe CW | Soil Low Fe.0 |
| 6 – 9 cm | Electrode 1 | High Fe CW | Soil High Fe.1 |
|  |  |  | Biof. High Fe.1 |
|  |  | Low Fe CW | Soil Low Fe.1 |
|  |  |  | Biof. Low Fe.1 |
| 12 – 15 cm | Electrode 2 | High Fe CW | Soil High Fe.2 |
|  |  |  | Biof. High Fe.2 |
|  |  | Low Fe CW | Soil Low Fe.2 |
|  |  |  | Biof. Low Fe.2 |
| 18 – 21 cm | Electrode 3 | High Fe CW | Soil High Fe.3 |
|  |  |  | Biof. High Fe.3 |
|  |  | Low Fe CW | Soil Low Fe.3 |
|  |  |  | Biof. Low Fe.3 |
| 24 – 27 cm | Electrode 4 | High Fe CW | Soil High Fe.4 |
|  |  |  | Biof. High Fe.4 |
|  |  | Low Fe CW | Soil Low Fe.4 |
|  |  |  | Biof. Low Fe.4 |
| 30 – 33 cm | Electrode 5 | High Fe CW | Soil High Fe.5 |
|  |  |  | Biof. High Fe.5 |
|  |  | Low Fe CW | Soil Low Fe.5 |
|  |  |  | Biof. Low Fe.5 |

### DNA extraction, *Acidimicobiaceae* bacterium A6 quantification and phylogenic analysis

Total genomic DNA was extracted from each biofilm sample obtained in duplicate from a half face or full face of each electrode deployed in the field, except for the deep electrode of set 6, which was only partially recovered and only one sample could be obtained from all faces. Since many electrodes in the CWs had a much lower biofilm mass than the filed electrodes, only one sample was recovered for DNA extraction from the CW electrodes. DNA was also extracted from each soil sample as described above (Table S1). Extractions were done using the FastDNA^®^ spin kit for soil (MP Biomedicals, USA) according to the manufacturer’s instructions. Total DNA was eluted in 100 μl of sterile water and its concentrations were measured using Qubit 2.0^®^ (Invitrogen, USA). All DNA samples were preserved at -20 °C until further analysis.

Bacteria quantification was carried out via Quantitative PCR (qPCR) using the Applied Biosystems StepOnePlus^™^ Real Time PCR system. A6 quantification was done by amplifying a section of the 16s rRNA gene between the variable regions V1 and V4 using primer set 33F/232R (33F: 5’ -GGCGGCGTGCTTAACACAT-3’ / 232R: 5’-GAGCCCGTCCCAGAGTGATA-3’). *Geobacter* spp., an electrogenic bacteria known for its ability colonize electrodes, were quantified by amplifying a region of the 16s rRNA gene using primer set 561F/825R (561F: 5’-GCGTGTAGGCGGTTTCTTAA-3’ / 825R: 5’ATCTACGGATTTCACTCCTACA-3’).

Each qPCR mixture (20 μL) was composed of 10 μL of SYBR Premix Ex Taq^®^ II 2X (Takara, Japan), 0.8 μL of each forward and reverse 10 μM primer, and DNA template. Thermal cycling conditions were initiated for 30 s at 95 °C, followed by 40 cycles with varying times and temperature depending on the amplicon being generated, and ended with a melting curve analysis for SYBR Green assay used to distinguish the targeted PCR product from the non-targeted PCR products. For A6 amplification, the cycling consisted of 10 s at 95 °C, 15 s of annealing at 59 °C and 15 s at 72 °C. For total bacteria quantification, each cycle consisted of 5 s at 94 °C, 30 s at 55 °C and 30 s at 70 °C. Each qPCR reaction was run in duplicate or triplicate per sample and included negative controls and a standard curve; the last one consisting of serial dilutions of known numbers of copies of DNA of the gene per volume. Finally, the results were converted into copies of DNA / m^2^ by dividing the total gene copies obtained from qPCR by the surface area of sediments for soil samples or by the surface area of the electrode for the electrode biofilm samples.

In order to determine the microbial community composition and abundance in the sediments (field and CW) and compare it to that formed on the electrodes, sequencing and phylogenetic analysis was performed by Novogene (Beijing, China) as follows: From total genomic DNA, the variable region V4 of the 16s rRNA gene was amplified using the primer set 515F/806R (515 F: 5′-GTGCCAGCMGCCGCGGTAA-3′ / 806R: 5’ -GGACTACHVGGGTWTCTAAT-3’) with a barcode following the method of Caporaso *et al* (2011) . All PCR reactions were carried out with Phusion^®^ High-Fidelity PCR master mix (New England Biolabs). PCR products quantification and qualification were determined by electrophoresis on 2% agarose gel. The resulting amplicons were pooled, purified, quantified. Sequencing libraries were generated using TruSeq^®^ DNA PCR-free sample preparation kit (Illumina, USA) following the manufacturer’s protocol and index codes were added. The library quality was assessed on the Qubit@ 2.0 Fluorometer (Thermo Scientific) and Agilent Bioanalyzer 2100 system. Finally, sequencing was performed on an IlluminaHiSeq2500 platform and 250 bp paired-end reads were generated.

Paired-end reads were assembled by using FLASH V.1.2.7 (43). Raw reads were processed according to QIIME V1.7.0 quality controlled process (44) and chimeric sequences were filtered out using UCHIME algorithm (45). For all samples (field and CW) a minimum of 25,000 sequences were obtained. These resulting sequences were clustered into operational taxonomic units (OTUs) using Uparse V7.0.1001 (46). Sequences with ≥97% similarity were assigned to the same OTUs. A total of 3206 OTUs were produced across all field samples and 2870 OTUs were produced across all CW mesocosm samples, with a range between 1422-1794 OTUs per field sample and 674 – 1481 OTUs per mesocosm sample. A representative sequence for each OTU was screened for taxonomic annotation using the Ribosomal Database Project (RDP) Classifier (47, 48) using GreenGene database(49) at a minimum of 80% confidence threshold for all OTUs. For CW samples, the OTUs were screened for taxonomic annotation applying the blastn algorithm against the 2016 NCBI’s 16s ribosomal RNA sequences for bacteria and archaea at an e-value of 1e^−5^. A6’s 16s rRNA gene sequence was included to NCBI’s database for annotation at the family and genus level of the top 100 and 300 most abundant OTUs. Finally, samples were standardized using the least sequence number obtained from all samples so that the same number of sequences were used for calculating the relative abundance of OTUs.

### Soil surface area analysis

Nitrogen sorption was used to determine the surface area of the soil samples taken from the field (Table 2) and the CW (Table 3). Prior to the analysis, samples were oven-dried at 56°C until the mass stabilized. Subsequently, the samples were degassed at 60°C and 0.1 mmHg using a Smart Micrometrics VacPrep (Norcross, GA, USA). The nitrogen sorption measurements were conducted using a Micromeritics 3FLEX (Norcross, GA, USA), using the BET method (Brunauer–Emmett–Teller) to calculate the surface area of the soil. The measurements obtained were used to normalize the bacterial count data by surface area.

### Analytical Methods

The sediment’s iron concentrations were analyzed using the ferrozine method (50). Briefly, 0.5 mL sediment sample was added to 9.5 mL 0.5 M HCl and shaken for 24 hours at room temperature to extract Fe(II). In total, 60 μL 6.25 M NH_2_OH·HCl was added to 3 mL of extraction solution and shaken for 24 hours at room temperature to reduce Fe(III) to Fe(II). For the chromogenic reaction, 60 μL of extraction solution was added to 3 mL 1 g/L ferrozine solution (pH 7.0) and reacted for 30 minutes. The concentrations of Fe(II) and total Fe were measured by reading the absorbance at the 562-nm wavelength using a Spectronic^®^ Genesys^™^ 2 instrument, and the Fe(III) concentration was calculated from the difference. Oxidation-reduction potential (ORP) at different depths of CW mesocosms were measured using probes from Thermo Scientific, Inc. During the operation of CW mesocosms, electrode pairs were only connected using wires, whereas 1000-Ω resistors were connected in the circuit for voltage measurement. The voltages between electrode pairs in the CW mesocosms were measured using a multimeter and currents were calculated accordingly.

### Statistical Analysis

The Welch t-test statistical analysis was used to determine if there were statistical differences in bacterial counts per surface area between electrode and soil samples in the field and CW mesocosm experiments. Paired two sample t-tests were conducted to determine if there were statistical differences in relative abundance for electrode/soil pairs in the CW mesocosms.

## ACKNOWLEDGEMENTS

Funding for this research were provided by the Program of International S&T Cooperation, National Key Research and Development Program of China (2016YFE0106600). The Instituto de Fomento al Talento Humano – Ecuador and the Mary and Randall Hack ’69 Research Grant provided funding for Melany Ruiz-Urigüen. We thank Dr. Claire White and Dr. Nishant Garg for their collaboration in soil surface area analysis.

---

**Figure 1.**
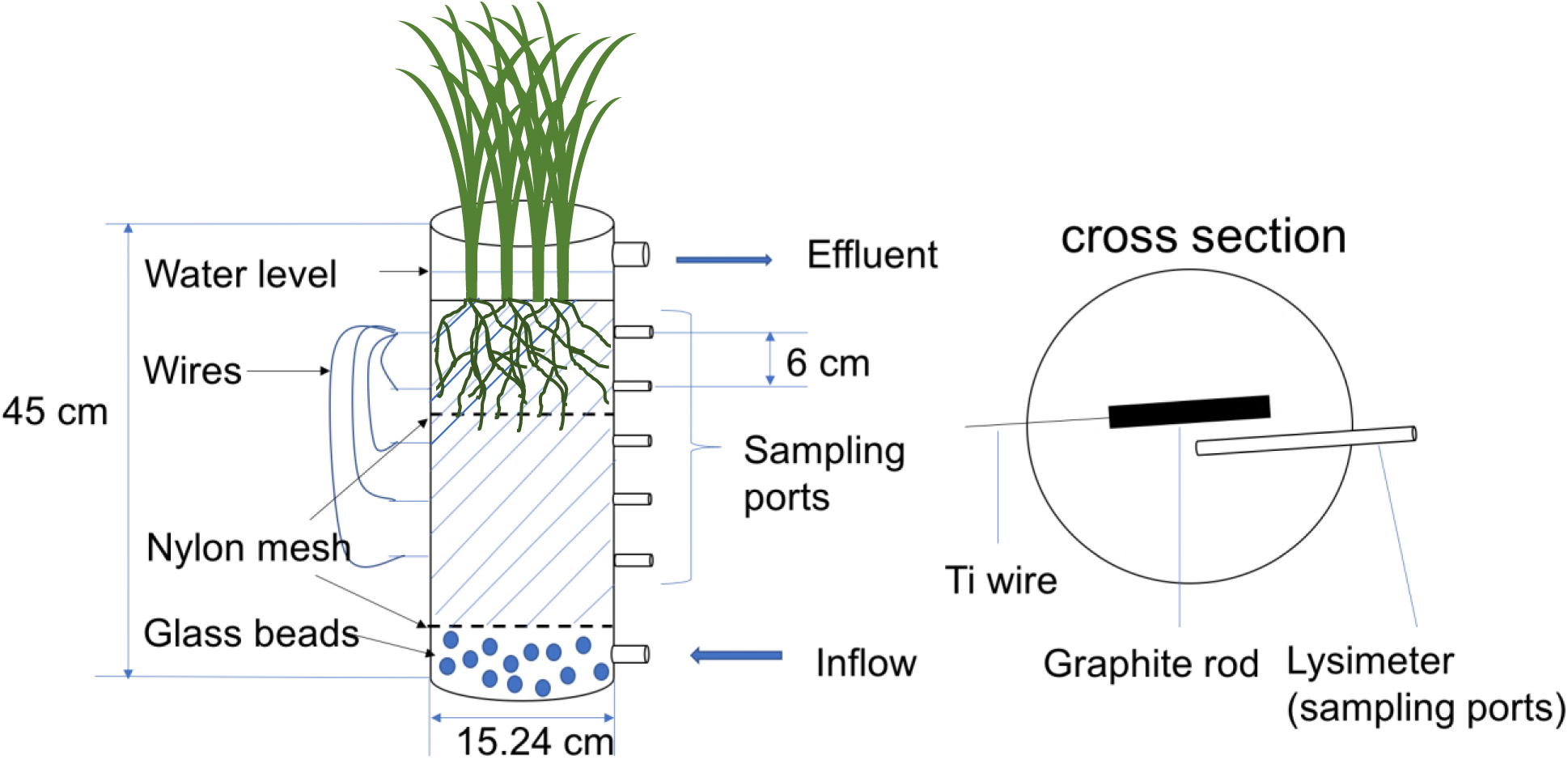
Schematic of a CW mesocosm and electrode setup.

